# The Cortical Spectrum: a robust structural continuum in primate cerebral cortex revealed by histological staining and magnetic resonance imaging

**DOI:** 10.1101/2021.09.09.459678

**Authors:** Yohan J. John, Basilis Zikopoulos, Miguel Ángel García-Cabezas, Helen Barbas

## Abstract

High-level characterizations of the primate cerebral cortex sit between two extremes: on one end the cortical mantle is seen as a mosaic of structurally and functionally unique areas, and on the other it is seen as a uniform six-layered structure in which functional differences are defined solely by extrinsic connections. Neither of these extremes captures the crucial neuroanatomical finding: that the cortex exhibits systematic gradations in architectonic structure. These gradations have been shown to predict cortico-cortical connectivity, which in turn suggests powerful ways to ground connectomics in anatomical structure, and by extension cortical function. A challenge to more widespread use of this concept is the labor-intensive and invasive nature of histological staining, which is the primary means of recognizing anatomical gradations. Here we show that a novel computational analysis technique can be used to derive a coarse-grained picture of cortical variation. For each of 78 cortical areas spanning the entire cortical mantle of the rhesus macaque, we created a high dimensional set of anatomical features derived from captured images of cortical tissue stained for myelin and SMI-32. The method involved semi-automated de-noising of images, and enabled comparison of brain areas without hand-labeling of features such as layer boundaries. We applied nonmetric multidimensional scaling (NMDS) to the dataset to visualize similarity among cortical areas. This analysis shows a systematic variation between weakly laminated (limbic) cortices and sharply laminated (eulaminate) cortices. We call this smooth continuum the ‘cortical spectrum’. We also show that this spectrum is visible within subsystems of the cortex: the occipital, parietal, temporal, motor, prefrontal, and insular cortices. We compared the NMDS-derived spectrum with a spectrum produced using T1- and T2-weighted magnetic resonance imaging (MRI) data derived from macaque, and found close agreement of the two coarse-graining methods. This evidence suggests that T1/T2 data, routinely obtained in human MRI studies, can be used as an effective proxy for data derived from high-resolution histological methods. More generally, this approach shows that the cortical spectrum is robust to the specific method used to compare cortical areas, and is therefore a powerful tool to understand the principles of organization of the primate cortex.

## 1. Introduction

Among descriptions of cortical architecture, two diametrically opposed pictures can be posited. At one extreme, the cortex is presented as a patchwork quilt of distinct areas separated by defined boundaries. The century-old architectonic map of Korbinian Brodmann can create this impression: the order of area numbers is arbitrary, conveying no information about similarities and differences of architectonic structure (Brodmann, 1909). At the other extreme, the entire cortex is presented as a uniform structure (Rockel et al., 1980; Carlo & Stevens, 2013), which is often described in terms of canonical columns or microcolumns (reviewed in Horton & Adams, 2005). This perspective can create the impression that cortical areas are equivalent, modular computational units, and that differences in function arise solely from differences in extrinsic connectivity. Both pictures of cortical structure are misleading. Further, using cytoarchitectonic features to divide the cortex into discrete areas has not led to consensus: researchers have not agreed on consistent criteria for drawing areal boundaries, which are often subtle (A. W. Campbell, 1905; Brodmann, 1909; von Economo et al., 1925; von Economo, 1927).

On the other hand, evidence that cortical architecture is not uniform across areas (Collins et al., 2010) has provided the basis to observe that the cellular and laminar features of areas vary systematically, so that each area is not totally discontinuous from its neighbors (reviewed in Sanides, 1970; Barbas, 2015; García-Cabezas et al., 2019, 2020). Following the approach of von Economo and collaborators (von Economo, 1927; von Economo et al., 1925), investigators began classifying cortical areas according to broad types, based on laminar features that vary in a graded manner, such as the number and prominence of layers and the sharpness of transitions between layers. Here we refer to this gradation of the cortical layering as the ‘degree of lamination’.

Several structural features have now been shown to co-vary with the gradient of laminar elaboration (Pandya et al., 1988; Huntenburg et al., 2018; Palomero-Gallagher & Zilles, 2018). Cortical types must be understood as discretizations of these continuous gradients. The number of cortical types depends on pragmatic considerations, and can vary from three (Zhang et al., 2020) to as many as eight (Hilgetag et al., 2016), but their ordering, based on degree of lamination, and their topological arrangement are not arbitrary, allowing for comparison among typologies.

The one-dimensional gradient of degree of lamination has proven to be an accurate indicator of cortico-cortical connectivity, enabling prediction of the pattern of laminar projections from one area to another. The linkage of connections to the systematic variation of the cortex, known as the Structural Model (Barbas, 1986; Barbas & Rempel-Clower, 1997), has been validated for all cortico-cortical areas thus far studied, and offers a unified framework for studying cortical structure, function, development, and evolutionary origin (reviewed in Barbas, 2015; García-Cabezas et al., 2019).

Determining cortical type by examination of Nissl-stained sections (García-Cabezas et al., 2020) requires training in neuroanatomy, and retains an element of subjectivity. To avoid these challenges, quantitative proxies for degree of lamination have been used, such as overall neural density (Dombrowski et al., 2001; Cahalane et al., 2012; Hilgetag et al., 2016). However, estimating neural density is a labor-intensive process that has not been automated yet, and it is not always the best proxy for cortical type. It is therefore necessary to develop coarse-grained estimates of cortical structure that can be used by researchers without extensive training in neuroanatomy. A coarse-grained synopsis of cortical structure can provide a bird’s eye view that is essential for identifying systematic structural patterns. Further, as new histological and gene expression approaches are introduced (Herculano-Houzel et al., 2013; Palomero-Gallagher & Zilles, 2018), they can be incorporated into this synoptic method.

Here we used a novel computational procedure applied to analysis of photomicrographs of cortical areas using cellular and axonal architectonic markers. From these photomicrographs we computed a dataset of cortical profiles of 78 areas in the rhesus macaque brain. To reduce the dimensionality of this dataset for the purposes of visualization and interpretation, we performed non-metric multidimensional scaling (NMDS). This enabled an integrated view of structural similarities and differences among cortical areas. This approach allowed visualization of the systematic gradation in structure across the entire cortex.

We also showed that the same gradation, which we call the ‘cortical spectrum’, is reflected in magnetic resonance imaging (MRI) data, which are collected broadly in humans. Specifically, we contrasted our architectonic results with analysis of rhesus macaque MRI data. For humans, MRI-based methods are now the most widespread means of determining anatomical structure (Glasser et al., 2016). We show that T1- and T2-weighted MRI signals provide qualitative results that are in agreement with the more high-resolution analysis based on stained tissue. This raised the possibility that MRI analysis, which can be challenging to interpret, may serve as a proxy for architectonic analysis, and more specifically for inferring the degree of lamination in the cortex in humans using non-invasive methods.

## 2. Methods

We used coronal brain sections from two rhesus macaque monkeys (cases AN and AQ) to analyze a total of 78 cortical regions, from one hemisphere in each case. These areas span the entire cortex, but are not an exhaustive sample of structural gradation, so the smooth gradation in structure is not fully reflected for every region. These cases have been used in other unrelated studies (e.g., García-Cabezas et al., 2017; Zikopoulos et al., 2018), so here we will describe a brief overview of tissue processing. Perfusion and sectioning of the brains were performed as reported previously (Barbas & Rempel-Clower, 1997; Zikopoulos et al., 2018). Experiments were performed in accordance with the National Institutes of Health Guide for the care and use of laboratory animals (publication 80–22 revised, 1996). Protocols were approved by the Institutional Animal Care and Use Committee at Boston University School of Medicine, Harvard Medical School, and New England Primate Research Center. Tissue sections were 40 micrometers (μm) thick. For each case, one set of sections was stained for myelin (Gallyas), and a matched set was stained for SMI-32, an antibody to a neurofilament protein that labels a subset of pyramidal projection neurons in layers II-III and V-VI (M. J. Campbell & Morrison, 1989; Hof et al., 1995). Sections were photographed at 40x magnification. These photomicrographs were then rotated and cropped to highlight a segment extending from the pia to the superficial white matter. Figure 1 shows examples of such photomicrographs for two cortical systems in the brain that include areas in motor and occipital cortices. The vertical height of each photograph varied depending on the cortical thickness. Widths also varied, as cropping was constrained by the amount of cortical curvature. Each cropped color image was first converted to grayscale using Matlab’s ‘rgb2gray’ function, and then inverted: each pixel’s gray value was subtracted from 255 to create a gray level. Thus, the pixel values varied in proportion to the darkness of the stain.

**Figure 1.**
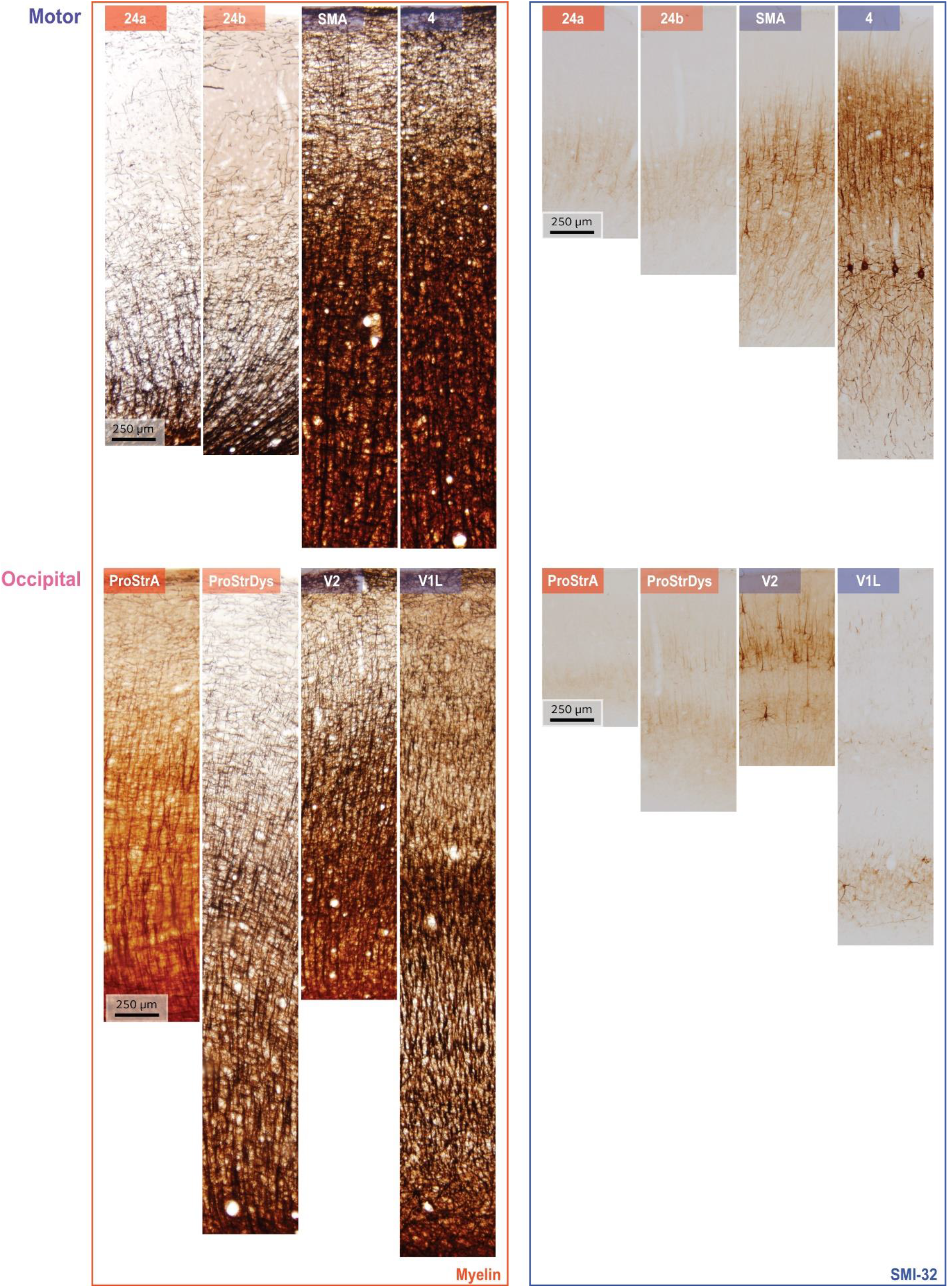
Examples of myelin staining (left column) and SMI-32 staining (right column). Rows show data from two systems, arranged in order from limbic to eulaminate. **Top:** areas along the motor cortical axis, **bottom:** areas along the occipital cortical axis. Variations in cortical thickness from pia to white matter arise due to non-uniform shrinkage during tissue processing.

A key challenge with histological staining methods is variability of baseline density of staining from case to case, and also of overall staining across sections within a case. In addition, staining artifacts can arise during experiments. Classically, when such sections are analyzed, the noise and variability are filtered out through subjective discernment of the relevant qualitative features. This skilled filtering has proved difficult to automate with machine learning, given the absence of vast hand-labeled datasets. For myelin-stained sections, the key qualitative features include the density of labeled myelin fibers, the presence and spacing of fiber bands, and the extent of laminar penetration of fibers. For SMI-32-stained sections, the qualitative features include the presence, size and laminar specificity of labeled neurons. Qualitative analyses, which are necessary for subsequent quantitative stereological estimation, are impractical if the goal is to study all cortical areas together. We therefore developed a semi-automated method to extract features from a large number of cortical areas (Figure 2). We first processed each image to generate binarized “all-or-nothing” labeled pixels (Figure 2B, E). We determined a threshold for each stained image: pixels above the threshold were assigned a non-zero value of unity. To estimate these thresholds in an unbiased fashion, we presented the images to three researchers in pseudorandom order, with the area labels withheld. We instructed the researchers to adjust the thresholds in order to balance a trade-off between the architectonic features and the noise, background, and staining artifacts. We designed custom software for this purpose, in which the original image was shown side-by-side with the thresholded binarized image. The researchers were instructed to step through the images and use a slider to adjust the threshold for each. For a given cortical area in a case, the threshold for a given stain type, myelin or SMI-32, was averaged from the estimates made by the researchers. The binarized images were used to compute cortical profiles. The researchers’ thresholds showed close agreement.

**Figure 2.**
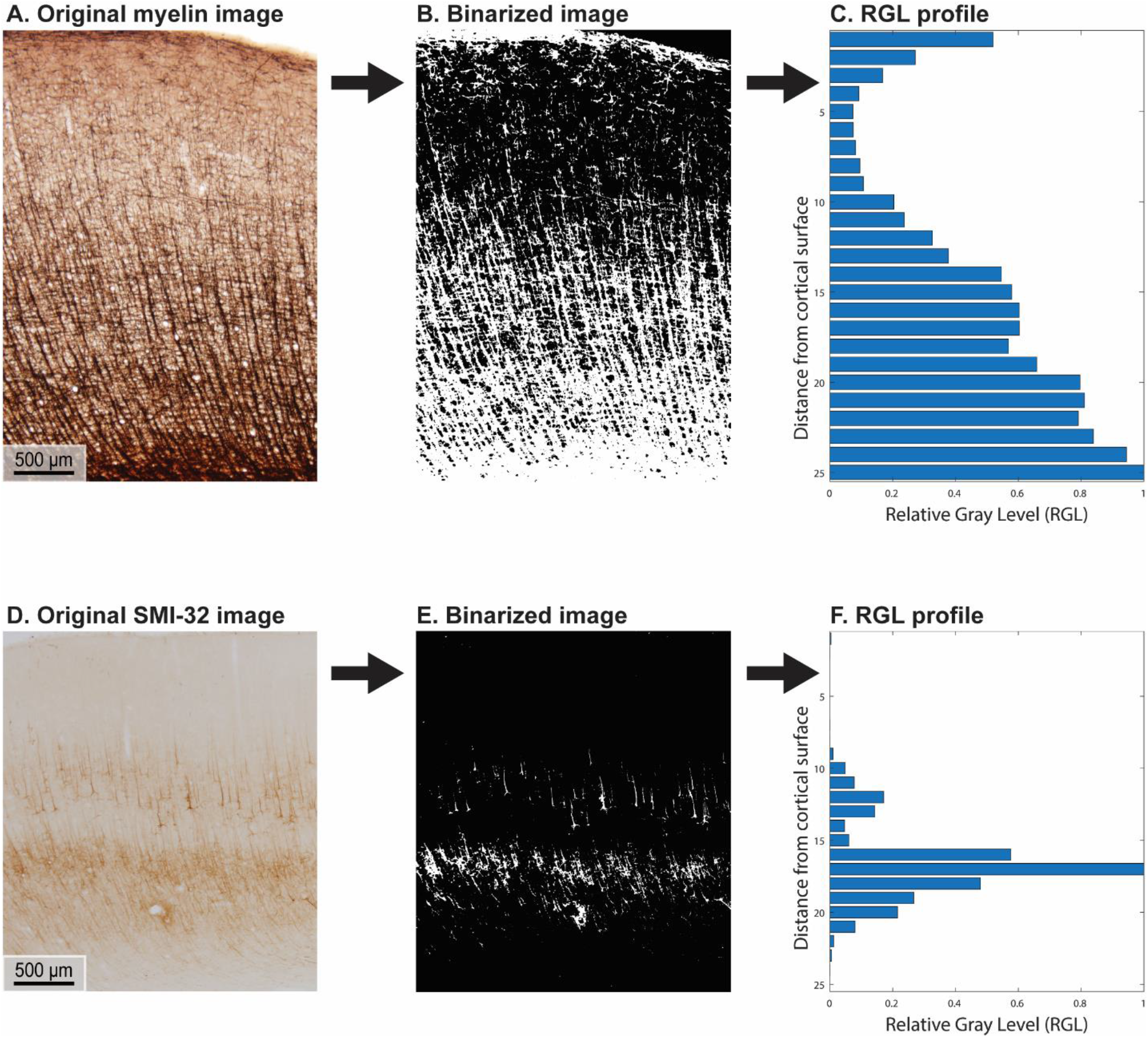
Example of image processing used to create the feature vector for each cortical area. A. Original photomicrograph of myelin stain of area 46 dorsal. B. Corresponding binarized image. C. Relative gray level (RGL) profile generated from the binarized image. D-F: analogous steps for a photomicrograph of SMI-32 stain of area 46 dorsal.

For each binarized image, we computed a cortical profile by dividing the image into equal-sized sub-images along the direction normal to the surface (the pia-to-white-matter direction) and averaging the pixel intensity within each sub-image or bin. These profiles were then normalized by the largest bin value. This resulted in a relative gray level (RGL) profile, which captures the relative variation of the stain with depth from the cortical surface (Figure 2C, F). Analyses were conducted for binarized images cropped to be of equal width, as well as for the uncropped binarized images. Results were qualitatively similar in both cases, so the results shown here are from the analysis of the uncropped images. We divided each image into 25 bins, which exceeds the standard number of layers commonly used to analyze cortical structure. Figure 2 shows an example of the image processing stages for area 46 dorsal (case AN).

The 25-dimensional RGL profiles from the myelin and SMI-32 images were concatenated to create a 50- dimensional feature vector for each cortical area. Each of the 78 feature vectors was the result of averaging feature vectors across the two cases. This averaged feature set was used to create a distance matrix using the L2 norm, which served as the basis for non-metric multidimensional scaling (NMDS). This method enables high-dimensional data points to be mapped into a lower dimensional space that preserves a nonlinear but monotonic transform of the distances among the points. A small number of dimensions, three in this case, facilitates visualization of the differences among cortical areas. The feature set was z-scored and then used to create a distance matrix among the 78 brain areas. For the comparison of different coarse-grained methods, we also computed the average RGL values for each binarized image. For comparison, cortical areas were classified by experts (BZ and MAG-C) using matched Nissl-stained sections, in accordance with previously published criteria (Pandya et al., 1988; García-Cabezas et al., 2020).

Image rotation and cropping was performed using ImageJ. Further processing and statistical analyses were performed using MATLAB. NMDS was performed using MATLAB’s in-built function.

For the analysis of MRI data, we used the T1-weighted (T1w) and T2-weighted (T2w) datasets of a shared rhesus macaque MRI scan from McGill University obtained from the NeuroImaging Tools & Resources Collaboratory (www.nitrc.org), and we estimated the T1w/T2w ratio. The dataset is available under the Creative Commons – Attribution-Non-Commercial Share Alike (CC-BY-NC-SA)-Standard INDI data sharing policy, which prohibits use of the data for commercial purposes. The T1w/T2w ratio reflects the content of intracortical myelin, and is widely used in imaging studies of cortical hierarchies and connectivity (Glasser et al., 2016; Zhang et al., 2020).

## 3. Results

We used sections stained for myelin- and SMI-32- to capture and analyze photomicrographs from two rhesus macaque brains to create a combined dataset consisting of the myelin and SMI-32 profiles of 78 cortical areas. We used this high-dimensional dataset as the basis for NMDS analysis, which allowed us to transform high-dimensional cortical data into a three-dimensional space that preserved the relative distances among cortical areas. NMDS resulted in an acceptably low stress value (0.09), suggesting a good agreement between the high-dimensional dataset and the derived three-dimensional space. This allowed us to visualize the relationships of similarity and difference among the cortical areas. Scatter plots of the cortical areas in the NMDS-space showed a clear gradation of cortical areas: the less sharply laminated limbic areas were at one end of the continuum, and the more sharply laminated primary sensory and motor areas were at the other end (Figures 3 and 4). These results show that the cortex can be described as a structural spectrum.

**Figure 3.**
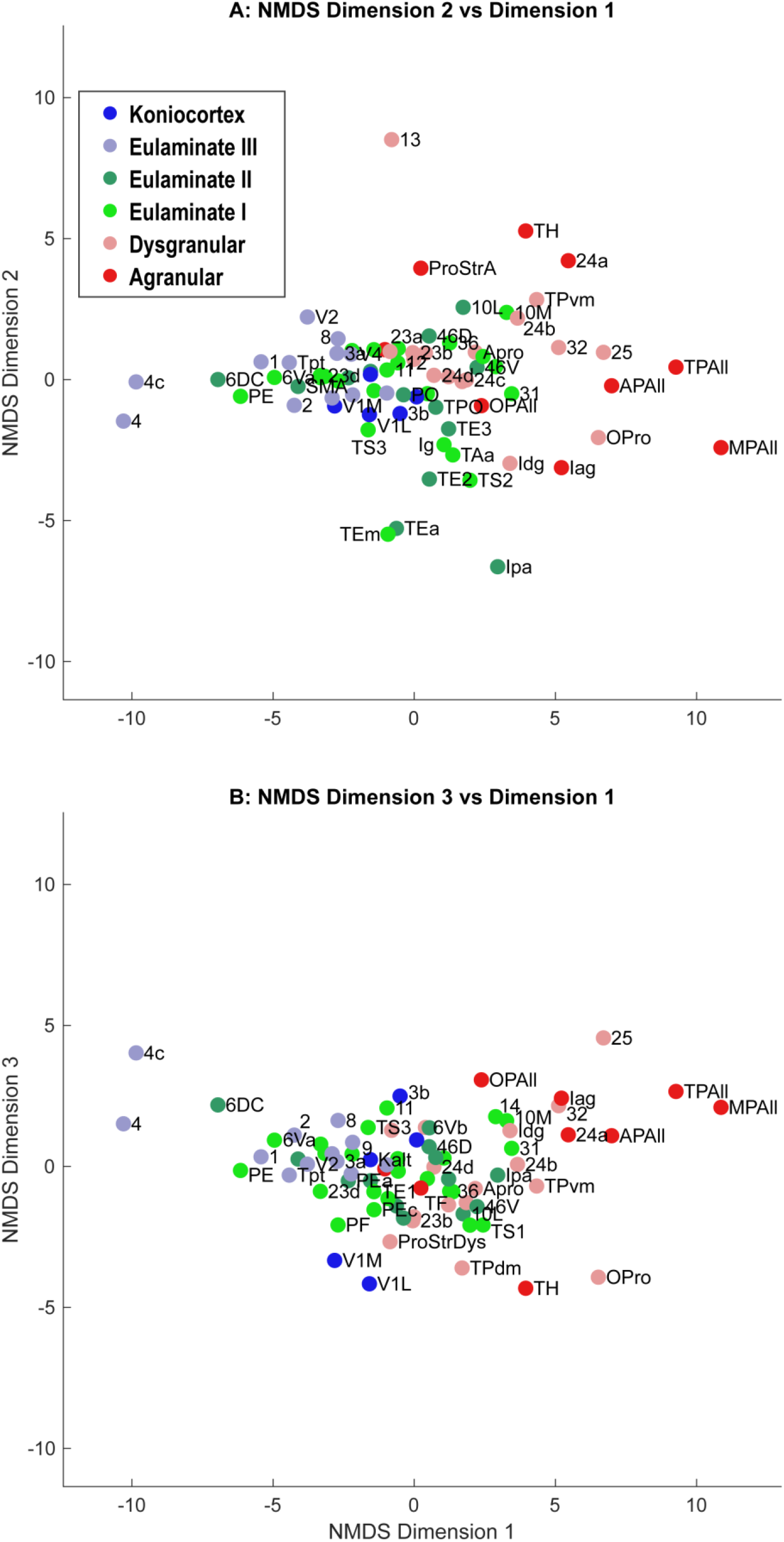
Scatter plot of 78 cortical areas in NMDS-derived dimensions. We used three dimensions for NMDS, which yielded a stress value of 0.09. Dimension 1 showed the clearest alignment with the subjective classification of cortical types according to degree of lamination. **A**. Dimension 2 versus Dimension 1. **B**. Dimension 3 versus Dimension 1. Due to partial overlap of data points, some area labels were omitted for clarity.

**Figure 4.**
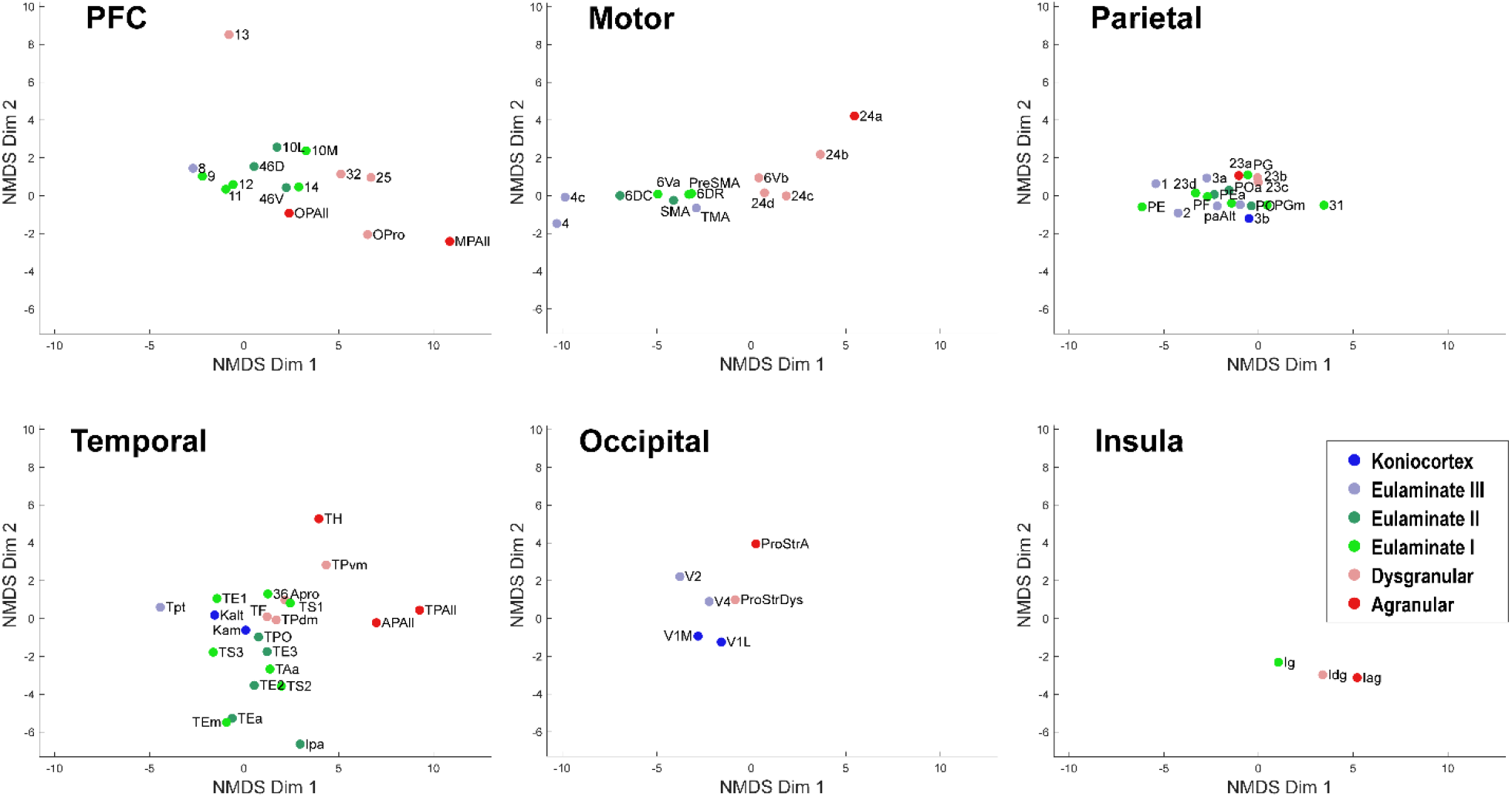
Scatter plot of 78 cortical areas in NMDS-derived dimensions, divided by cortical system. Only the first two dimensions are plotted. **A**. Prefrontal cortex, **B**. Motor cortex, **C**. Parietal cortex, **D**. Temporal cortex, **E**. Occipital cortex, **F**. Insular cortex. Due to overlap of data points, area labels were omitted from some of the points.

In Figures 3 and 4, the cortical areas were shown using color-coded text that represents the subjective categorization by experts based on previously published criteria (Pandya et al., 1988; García-Cabezas et al., 2020). This categorization is overlaid on a map of the cortex in Figure 5. We divided the 78 cortical areas into 6 discrete levels of lamination: agranular (red), dysgranular (pink), eulaminate I (light green), eulaminate II (dark green), eulaminate III (light blue), and koniocortex (dark blue). Agranular (lacking a granular layer IV) and dysgranular (having a weak or incipient layer IV) are collectively referred to as limbic cortices. The non-limbic cortices are collectively described as eulaminate (“well-laminated”), and are characterized by a visible layer IV. They show increasingly clear differentiation between layers II and III and between layers V and VI. Among eulaminate areas, koniocortex includes the most sharply laminated cortices, only seen in sensory areas. The primary motor cortex, as a specialized efferent system, has a distinctive layer V, and a small layer IV, but is strongly myelinated, like the well-laminated sensory association cortices (reviewed in García‐Cabezas & Barbas, 2014). Dimension 1 of the NMDS was found to reflect the discrete levels of lamination (Figure 2A, B). The limbic cortices were clustered near one end of Dimension 1 and the eulaminate cortices were clustered near the opposite end.

**Figure 5.**
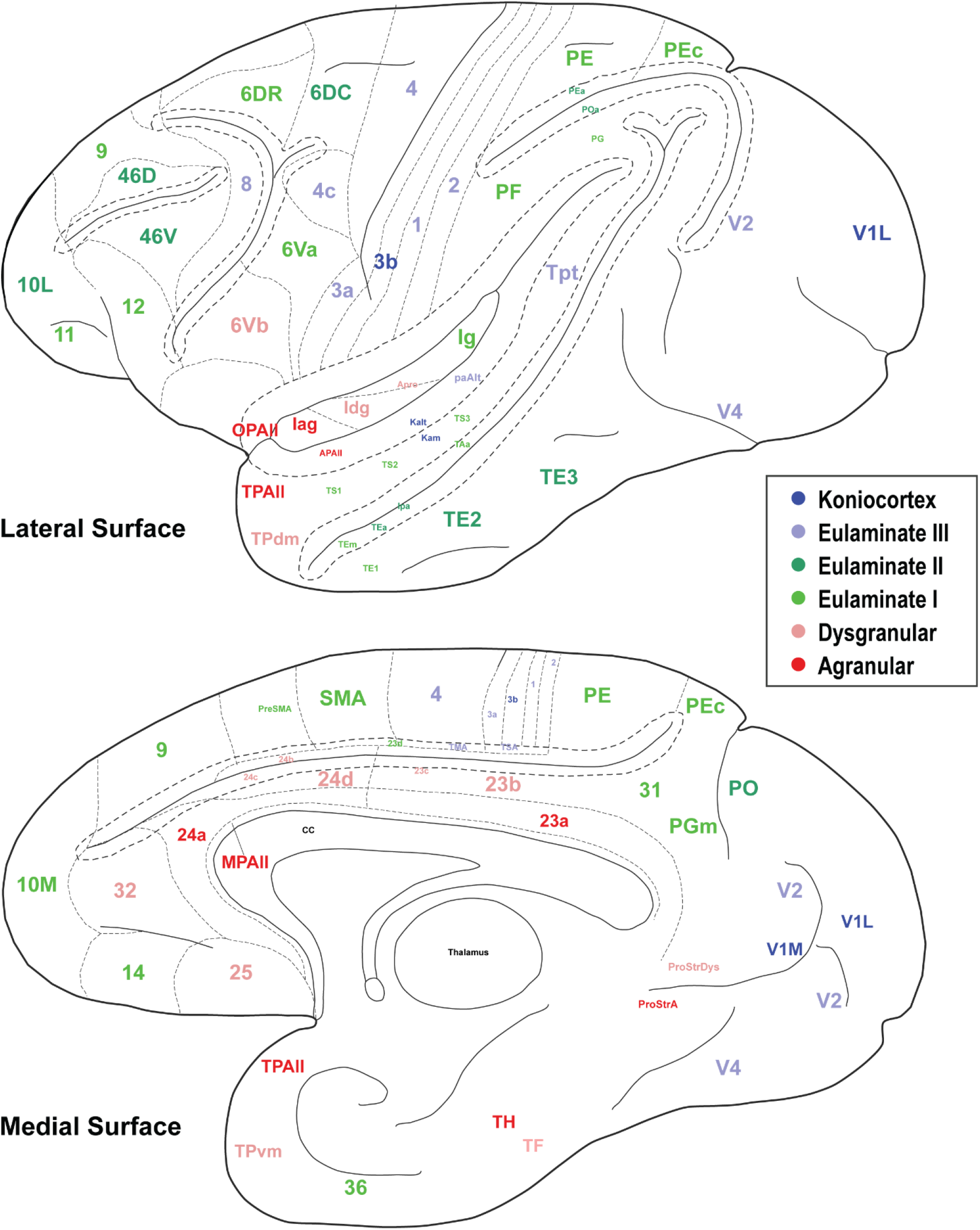
The cortical spectrum. 78 brain areas used in the analyses here. Colors correspond to the subjective categorization performed by experts. Title location reflects approximate center of region.

We plotted the same NMDS results according to cortical lobe or sub-system, which highlighted the fact that each sub-system contained its own structural spectrum (Figure 4). The prefrontal, motor, parietal, temporal, occipital and insular sub-systems each showed a spectrum or gradient that aligned with the degree of lamination, which corresponded primarily with Dimension 1. The motor cortex spectrum aligned most closely with the assigned levels, followed by the temporal cortex spectrum. The other sub-systems showed more mixing or ambiguity among cortical areas, while preserving a general limbic-to-eulaminate gradation. Interestingly, while the occipital limbic-to-eulaminate gradation is reflected in Dimension 1, it is more clearly visible along Dimension 2. The dimensions of NMDS are not straightforward to interpret, but they may suggest hypotheses for future research.

We plotted the T1- and T2-weighted voxel values for each of the 78 cortical areas analyzed above, resulting in analogous scatter plots (Figures 6 and 7). Due to the MRI resolution, each cortical area produced on average a single voxel of data per hemisphere. We averaged the data from the two hemispheres to produce a single two-dimensional dataset. As seen in Figure 6, the MRI data capture the limbic-to-eulaminate trend seen in Dimension 1 of the NMDS analysis. Specifically, the T1-weighted data is most informative of the degree of lamination. To facilitate further comparison with the NMDS method, we also plotted the MRI data according to sub-system in Figure 7, which can be contrasted with Figure 4.

**Figure 6.**
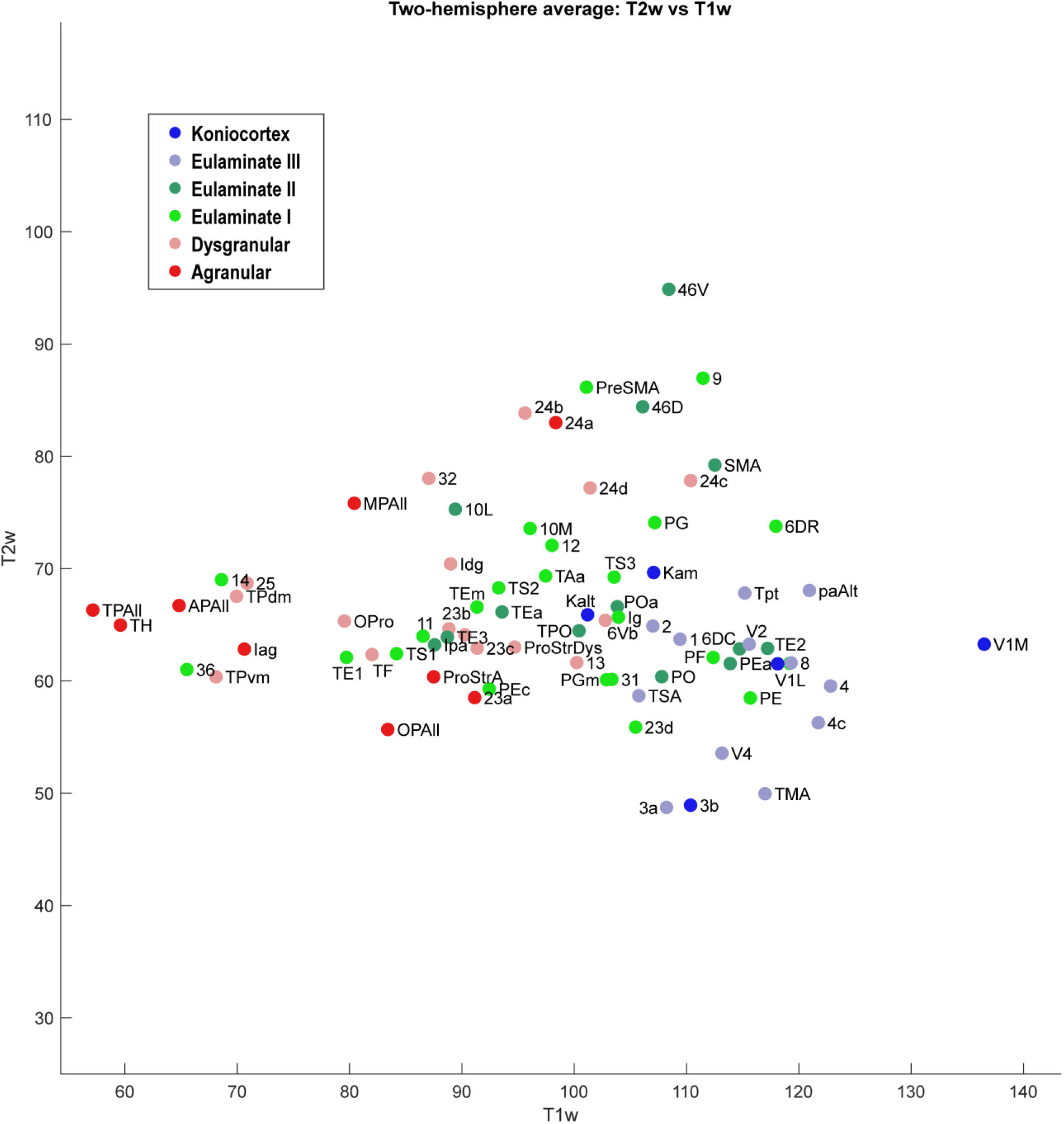
Scatter plot of 78 cortical areas. X-axis is T1-weighted MRI data, and Y-axis is T2-weighted MRI data. Each data point is the average of the optical density value for each area from the left and right hemisphere of the case analyzed.

**Figure 7.**
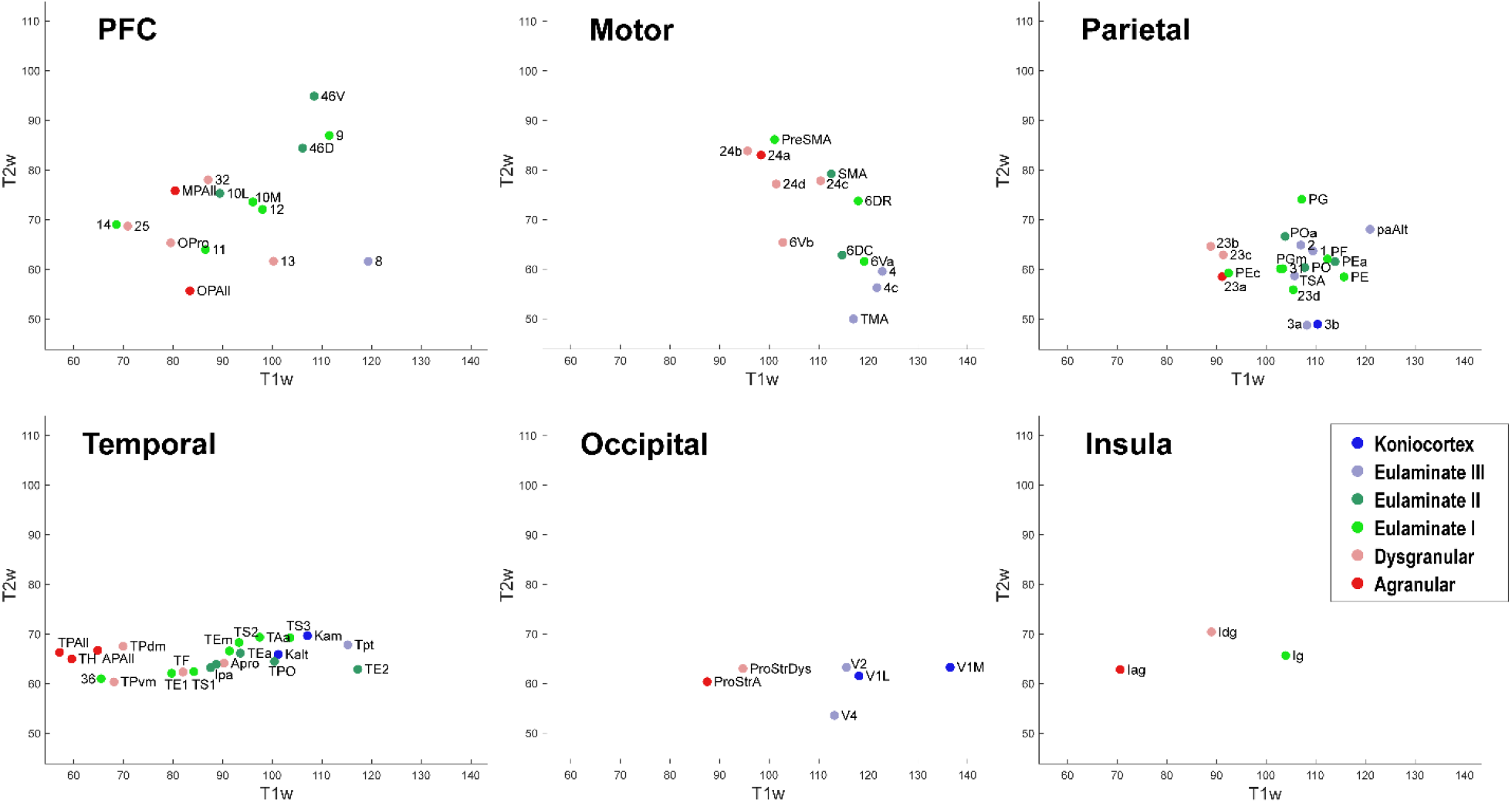
Scatter plot of T1w and T2w MRI data from 78 cortical areas, divided by system. **A**. Prefrontal cortex, **B**. Motor cortex, **C**. Parietal cortex, **D**. Temporal cortex, **E**. Occipital cortex, **F**. Insular cortex. Due to overlap of data points, area labels were omitted from some of the points for clarity.

To assess the quality of analyses based on cortical profiles and MRI, we grouped the cortices by the expert-labeled cortical type and plotted the corresponding z-scored values of mean relative gray level (RGL) in SMI-32 images (Figure 8A); mean RGL in Gallyas images (Figure 8B); the negative of Dimension 1 from the NMDS analysis of cortical profiles (Figure 8C); and T1-weighted MRI values (Figure 8D). Note that the NMDS dimensions are arbitrarily set by the algorithm, and in this case the limbic-to-eulaminate direction was anti-correlated with NMDS Dimension 1, so for comparison the sign was reversed. It can be seen that the cortical profile-based data (Figure 8C) and the T1-weighted MRI data both reflect the limbic-to-eulaminate spectrum.

**Figure 8.**
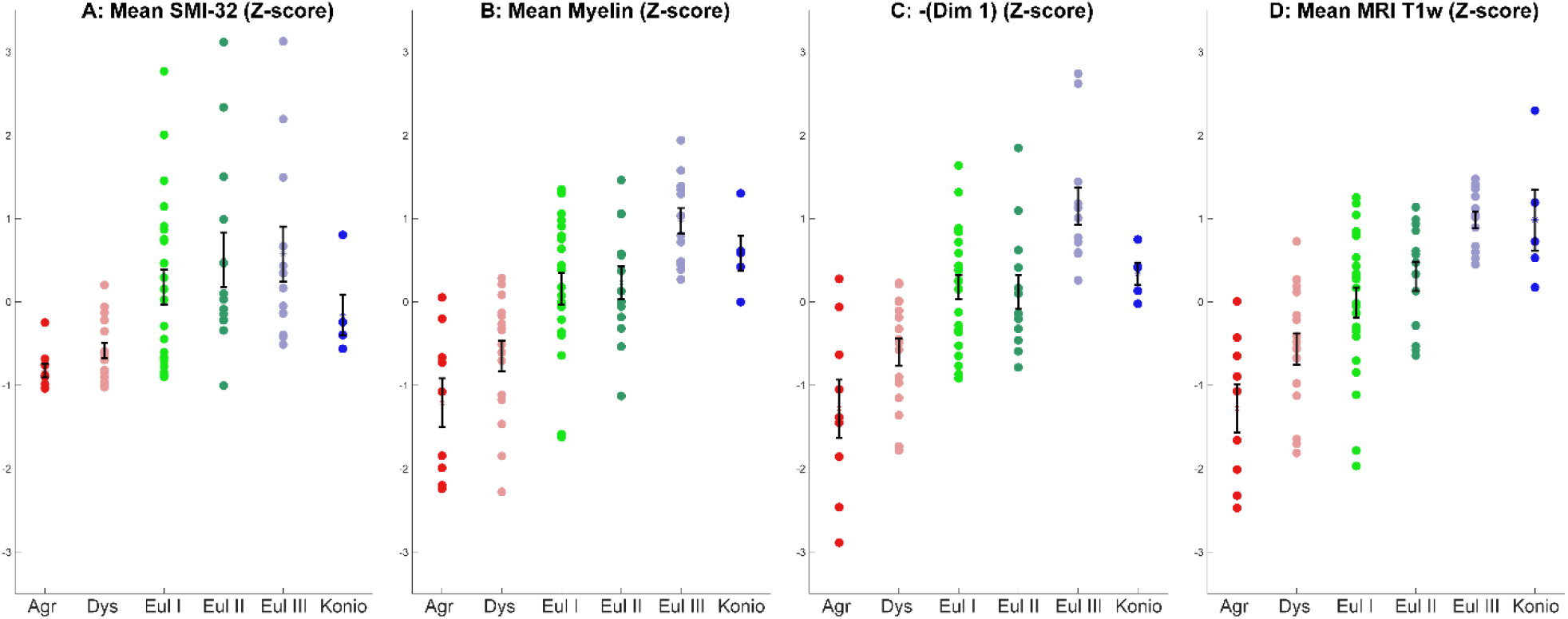
Comparison of four coarse-grained measurements that reflect the limbic-to-eulaminate spectrum. A. Mean SMI-32 RGL. B. Mean myelin RGL. C. Sign-flipped NMDS Dimension 1. D. T1-weighted data. All data are z-scored. Error bars show standard error of the mean.

## 4. Discussion

We showed that cortical profiles produced by analysis of cortical areas stained for myelin and SMI-32 support a conception of the cortical quilt as a structural continuum, which we refer to as the cortical spectrum. The spectrum derived from these cortical profiles shows broad agreement with cortical typologies derived from Nissl-stained cortical tissue, despite being performed without hand-labeling of layer boundaries. The Nissl stain has made it possible to identify cortical layers using transitions in features such as neuron arrangement and density. Once layer boundaries have been determined in cortical areas, the areas can be ordered according to the number of layers and the sharpness of transitions between layers, a pair of linked features which we refer to collectively as the degree of lamination. The degree of lamination is a key organizing principle of the cortex, as it enables prediction of the laminar pattern of origin and termination of cortico-cortical connections, their strength, and even presence or absence (Barbas, 1986; Barbas & Rempel-Clower, 1997; Hilgetag et al., 2016; Goulas et al., 2018; García-Cabezas et al., 2019). Quantitative estimates of the degree of lamination are therefore needed.

Neuron density, estimated from the widely used Nissl stain, serves as a partial proxy for degree of lamination, but is not suitable for all areas (García‐Cabezas & Barbas, 2014). Another example of a Nissl-derived estimate is ‘externopyramidalization’, which is the ratio of soma sizes of supragranular pyramidal neurons (above layer IV) to those of infragranular pyramidal neurons (below layer IV) that has been shown to correlate well with gradual cytoarchitectonic changes of the cortical mantle (Sanides, 1970; Pandya et al., 1988; Goulas et al., 2018; García-Cabezas et al., 2020). None of the quantitative features thus far identified is an ideal proxy for the assessment of an expert, but as such features are accumulated, the key concept becomes easier to recognize: cortical structure varies in a systematic and graded manner, resulting in a spectrum from limbic to eulaminate cortices. Figure 9 schematically represents how these feature gradients reflect degree of lamination.

**Figure 9.**
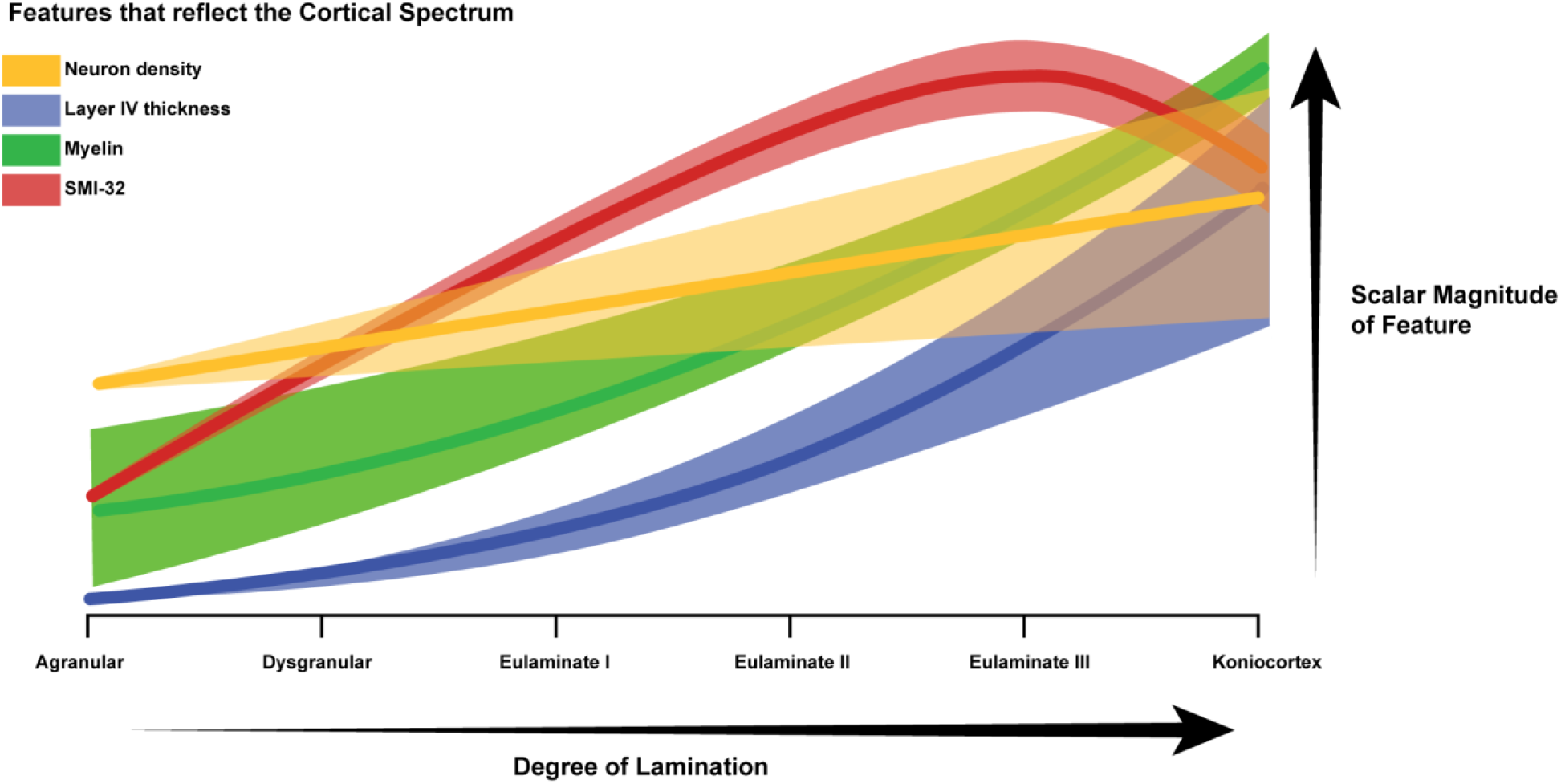
Schematic diagram of the cortical spectrum. The X-axis represents increasing degree of lamination, which corresponds to the traversal from agranular areas to koniocortical areas. The Y-axis represents the relative magnitudes of the features that serve as partial proxies of degree of lamination (arbitrary units). The thick translucent bar surrounding each plot corresponds to the variability in the feature. For example, the green bar corresponding to the variability in myelin density is initially thick and then becomes thinner, because a few agranular areas show high myelin, whereas eulaminate areas are always well laminated. Limbic areas generally have low myelin content, but cortical myelin shows not only the intrinsic myelin of an area but also terminations from pathways from other structures that may be myelinated.

Data derived from Nissl-stained tissue provide useful proxies for degree of lamination, but their estimation tends to be a labor-intensive process. For example, estimating neural density requires examining cortical slices under a microscope to identify and categorize cells to enable stereological estimation of the number and density of neurons or distinct cell types (Pakkenberg & Gundersen, 1997). For this reason, the Nissl-derived quantitative estimates of degree of lamination have only been used in specific sub-systems of the cortex. The method described here, by contrast, produces a relatively rapid bird’s eye perspective of structural variation across the entire cortical mantle. Moreover, the technique can be applied to existing databases of cortical photomicrographs, and can be scaled up for comparison across species as well as among individuals within a species.

Our results confirm and extend prior observations that myelin and SMI-32 reveal architectonic features that serve as partial proxies for the degree of lamination (García‐Cabezas & Barbas, 2014; García-Cabezas et al., 2017). In the case of myelin, the key feature is the extent to which myelin ‘reaches up’ from below layer VI and into the gray matter. Note that myelin labels both efferent and afferent pathways. Efferent pathways from limbic areas are the least myelinated among cortical areas, but afferent fibers to limbic areas vary in myelin content depending on origin. In the case of SMI-32, the key features are (1) the presence of labeling in the deep part of layer III, and (2) the presence of labeling in layers V and VI. The method presented here enables a computational estimation of these qualitative features, which can be discerned in normalized cortical profiles derived from photomicrographs of the stained tissue. Dimension 1 of the NMDS-space closely mirrors the subjectively-assigned cortical levels.

Importantly, our method does not involve manual parcellation of the cortex into layers, and therefore partly mitigates experimenter variability when deciding on criteria for layer boundaries. The broad agreement with Nissl-derived typologies can therefore be interpreted as providing support for the qualitative approach. Some bias can in principle arise during the image thresholding procedure that reduces noise and enhances signal, but we attempted to minimize this possibility by using a blind and randomized approach, and averaging across experimenters, who were not experts.

In humans, MRI is the primary means of studying structure noninvasively. Our results show that the cortical spectrum can also be discerned in T1-weighted MRI scans, which suggests a way to interpret this class of data. The large size of the human brain implies a higher-dimensional dataset than is possible in rhesus macaques for a given magnetic field strength, since multiple voxels will be available for each cortical area. Thus, the method outlined here – comparing cortical profiles using dimensionality reduction techniques such as NMDS -- may be well-suited to human MRI analyses. Given the widespread use of MRI, the cortical spectrum can be a powerful lens with which to contrast human brains in healthy and disordered subpopulations, and also to study individual differences.

In recent years, improvements in the resolution of MRI have enabled researchers to begin probing laminar structure, which is also described using terms such as microscale organization (van den Heuvel et al., 2015), cortical microstructure (Huntenburg et al., 2018), and microcircuits or microanatomy (Burt et al., 2018; reviewed in García-Cabezas et al., 2019). Neuroimaging data is already adding to the body of evidence in support of a central conclusion of the Structural Model: that structure predicts connectivity in the human brain (Zikopoulos et al., 2018; Paquola et al., 2020). The concept of a cortical spectrum can serve as a multi-level unifying principle, as it links low-dimensional structural gradients (Goulas et al., 2018; García-Cabezas et al., 2019, 2020; Goulas et al., 2021) with the higher-dimensional intricacies of laminar elaboration, which in turn predicts connectivity (Barbas, 1986; Barbas & Rempel-Clower, 1997), and therefore sheds light on functional interaction among cortical areas (Huntenburg et al., 2017; Vázquez-Rodríguez et al., 2019; Paquola et al., 2020). Moreover, the cortical spectrum generates testable hypotheses about neurodevelopment (Dombrowski et al., 2001; Barbas & García-Cabezas, 2016; García-Cabezas et al., 2019), which in recent years has become grounded in causal gradients of gene expression organizing factors (Puelles & Ferran, 2012; Puelles et al., 2019). Specifically, the continuous gradation of degree of lamination informs the search for the underlying genetic factors that bring about these gradations. In contrast to measurements such as thickness (Wagstyl et al., 2020) or ‘distance’ (Ercsey-Ravasz et al., 2013), this type of chemoarchitectonic patterning, which is reflected in the cortical spectrum, represents a more reliable lens with which to view cortical variation.

## Data and code availability

Data and code used in this study will be made available upon request.

## CRediT author statement

Conceptualization: YJ, BZ, MAG-C, HB; Formal analysis: YJ; Software: YJ; Funding acquisition: HB, BZ; Methodology: YJ, BZ, MAG-C; Writing: YJ, HB; Resources: HB, BZ.

## References

Barbas, H. (1986). Pattern in the laminar origin of corticocortical connections. The Journal of Comparative Neurology, 252(3), 415–422. https://doi.org/10.1002/cne.902520310

Barbas, H. (2015). General cortical and special prefrontal connections: Principles from structure to function. Annual Review of Neuroscience, 38, 269–289. https://doi.org/10.1146/annurev-neuro-071714-033936

Barbas, H., & García-Cabezas, M. Á. (2016). How the prefrontal executive got its stripes. Current Opinion in Neurobiology, 40, 125–134. https://doi.org/10.1016/j.conb.2016.07.003

Barbas, H., & Rempel-Clower, N. (1997). Cortical structure predicts the pattern of corticocortical connections. Cerebral Cortex (New York, N.Y.: 1991), 7(7), 635–646. https://doi.org/10.1093/cercor/7.7.635

Brodmann, K. (1909). Vergleichende Lokalisationslehre der Grosshirnrinde in ihren Prinzipien dargestellt auf Grund des Zellenbaues. Barth.

Burt, J. B., Demirtaş, M., Eckner, W. J., Navejar, N. M., Ji, J. L., Martin, W. J., Bernacchia, A., Anticevic, A., & Murray, J. D. (2018). Hierarchy of transcriptomic specialization across human cortex captured by structural neuroimaging topography. Nature Neuroscience, 21(9), 1251–1259. https://doi.org/10.1038/s41593-018-0195-0

Cahalane, D. J., Charvet, C. J., & Finlay, B. L. (2012). Systematic, balancing gradients in neuron density and number across the primate isocortex. Frontiers in Neuroanatomy, 6. https://doi.org/10.3389/fnana.2012.00028

Campbell, A. W. (1905). Histological studies on the localisation of cerebral function. University Press.

Campbell, M. J., & Morrison, J. H. (1989). Monoclonal antibody to neurofilament protein (SMI-32) labels a subpopulation of pyramidal neurons in the human and monkey neocortex. The Journal of Comparative Neurology, 282(2), 191–205. https://doi.org/10.1002/cne.902820204

Carlo, C. N., & Stevens, C. F. (2013). Structural uniformity of neocortex, revisited. Proceedings of the National Academy of Sciences, 110(4), 1488–1493. https://doi.org/10.1073/pnas.1221398110

Collins, C. E., Airey, D. C., Young, N. A., Leitch, D. B., & Kaas, J. H. (2010). Neuron densities vary across and within cortical areas in primates. Proceedings of the National Academy of Sciences, 107(36), 15927–15932. https://doi.org/10.1073/pnas.1010356107

Dombrowski, S. M., Hilgetag, C. C., & Barbas, H. (2001). Quantitative architecture distinguishes prefrontal cortical systems in the rhesus monkey. Cerebral Cortex (New York, N.Y.: 1991), 11(10), 975–988. https://doi.org/10.1093/cercor/11.10.975

Ercsey-Ravasz, M., Markov, N. T., Lamy, C., Van Essen, D. C., Knoblauch, K., Toroczkai, Z., & Kennedy, H. (2013). A Predictive Network Model of Cerebral Cortical Connectivity Based on a Distance Rule. Neuron, 80(1), 184–197. https://doi.org/10.1016/j.neuron.2013.07.036

García‐Cabezas, M. Á., & Barbas, H. (2014). Area 4 has layer IV in adult primates. European Journal of Neuroscience, 39(11), 1824–1834. https://doi.org/10.1111/ejn.12585

García-Cabezas, M. Á., Hacker, J. L., & Zikopoulos, B. (2020). A Protocol for Cortical Type Analysis of the Human Neocortex Applied on Histological Samples, the Atlas of Von Economo and Koskinas, and Magnetic Resonance Imaging. Frontiers in Neuroanatomy, 14. https://doi.org/10.3389/fnana.2020.576015

García-Cabezas, M. Á., Joyce, M. K. P., John, Y. J., Zikopoulos, B., & Barbas, H. (2017). Mirror trends of plasticity and stability indicators in primate prefrontal cortex. The European Journal of Neuroscience, 46(8), 2392–2405. https://doi.org/10.1111/ejn.13706

García-Cabezas, M. Á., Zikopoulos, B., & Barbas, H. (2019). The Structural Model: A theory linking connections, plasticity, pathology, development and evolution of the cerebral cortex. Brain Structure & Function, 224(3), 985–1008. https://doi.org/10.1007/s00429-019-01841-9

Glasser, M. F., Coalson, T. S., Robinson, E. C., Hacker, C. D., Harwell, J., Yacoub, E., Ugurbil, K., Andersson, J., Beckmann, C. F., Jenkinson, M., Smith, S. M., & Van Essen, D. C. (2016). A multi-modal parcellation of human cerebral cortex. Nature, 536(7615), 171–178. https://doi.org/10.1038/nature18933

Goulas, A., Changeux, J.-P., Wagstyl, K., Amunts, K., Palomero-Gallagher, N., & Hilgetag, C. C. (2021). The natural axis of transmitter receptor distribution in the human cerebral cortex. Proceedings of the National Academy of Sciences, 118(3). https://doi.org/10.1073/pnas.2020574118

Goulas, A., Zilles, K., & Hilgetag, C. C. (2018). Cortical Gradients and Laminar Projections in Mammals. Trends in Neurosciences, 41(11), 775–788. https://doi.org/10.1016/j.tins.2018.06.003

Herculano-Houzel, S., Watson, C. R., & Paxinos, G. (2013). Distribution of neurons in functional areas of the mouse cerebral cortex reveals quantitatively different cortical zones. Frontiers in Neuroanatomy, 7. https://doi.org/10.3389/fnana.2013.00035

Hilgetag, C. C., Medalla, M., Beul, S. F., & Barbas, H. (2016). The primate connectome in context: Principles of connections of the cortical visual system. NeuroImage, 134, 685–702. https://doi.org/10.1016/j.neuroimage.2016.04.017

Hof, P. R., Nimchinsky, E. A., & Morrison, J. H. (1995). Neurochemical phenotype of corticocortical connections in the macaque monkey: Quantitative analysis of a subset of neurofilament protein-immunoreactive projection neurons in frontal, parietal, temporal, and cingulate cortices. Journal of Comparative Neurology, 362(1), 109–133. https://doi.org/10.1002/cne.903620107

Horton, J. C., & Adams, D. L. (2005). The cortical column: A structure without a function. Philosophical Transactions of the Royal Society of London. Series B, Biological Sciences, 360(1456), 837–862. https://doi.org/10.1098/rstb.2005.1623

Huntenburg, J. M., Bazin, P.-L., Goulas, A., Tardif, C. L., Villringer, A., & Margulies, D. S. (2017). A Systematic Relationship Between Functional Connectivity and Intracortical Myelin in the Human Cerebral Cortex. Cerebral Cortex (New York, NY), 27(2), 981–997. https://doi.org/10.1093/cercor/bhx030

Huntenburg, J. M., Bazin, P.-L., & Margulies, D. S. (2018). Large-Scale Gradients in Human Cortical Organization. Trends in Cognitive Sciences, 22(1), 21–31. https://doi.org/10.1016/j.tics.2017.11.002

Pakkenberg, B., & Gundersen, H. J. (1997). Neocortical neuron number in humans: Effect of sex and age. The Journal of Comparative Neurology, 384(2), 312–320.

Palomero-Gallagher, N., & Zilles, K. (2018). Cyto-and receptor architectonic mapping of the human brain. Handbook of Clinical Neurology, 150, 355–387. https://doi.org/10.1016/B978-0-444-63639-3.00024-4

Pandya, D., Seltzer, B., & Barbas, H. (1988). Input-output organization of the primate cerebral cortex. Neurosciences, Comparative Primate Biology, 4, 39–80.

Paquola, C., Seidlitz, J., Benkarim, O., Royer, J., Klimes, P., Bethlehem, R. A. I., Larivière, S., Wael, R. V. de, Rodríguez-Cruces, R., Hall, J. A., Frauscher, B., Smallwood, J., & Bernhardt, B. C. (2020). A multi-scale cortical wiring space links cellular architecture and functional dynamics in the human brain. PLOS Biology, 18(11), e3000979. https://doi.org/10.1371/journal.pbio.3000979

Puelles, L., Alonso, A., García-Calero, E., & Martínez-de-la-Torre, M. (2019). Concentric ring topology of mammalian cortical sectors and relevance for patterning studies. Journal of Comparative Neurology, 527(10), 1731–1752. https://doi.org/10.1002/cne.24650

Puelles, L., & Ferran, J. L. (2012). Concept of neural genoarchitecture and its genomic fundament. Frontiers in Neuroanatomy, 6. https://doi.org/10.3389/fnana.2012.00047

Rockel, A. J., Hiorns, R. W., & Powell, T. P. (1980). The basic uniformity in structure of the neocortex. Brain: A Journal of Neurology, 103(2), 221–244. https://doi.org/10.1093/brain/103.2.221

Sanides, F. (1970). Functional architecture of motor and sensory cortices in primates in the light of a new concept of neocortex evolution. In The Primate Brain: Advances in Primatology (Noback CR, Montagna W, eds) (pp. 137–208).

van den Heuvel, M. P., Scholtens, L. H., Feldman Barrett, L., Hilgetag, C. C., & de Reus, M. A. (2015). Bridging Cytoarchitectonics and Connectomics in Human Cerebral Cortex. The Journal of Neuroscience: The Official Journal of the Society for Neuroscience, 35(41), 13943–13948. https://doi.org/10.1523/JNEUROSCI.2630-15.2015

Vázquez-Rodríguez, B., Suárez, L. E., Markello, R. D., Shafiei, G., Paquola, C., Hagmann, P., van den Heuvel, M. P., Bernhardt, B. C., Spreng, R. N., & Misic, B. (2019). Gradients of structure-function tethering across neocortex. Proceedings of the National Academy of Sciences of the United States of America, 116(42), 21219–21227. https://doi.org/10.1073/pnas.1903403116

von Economo, C. (1927). Zellaufbau der Grosshirnrinde des Menschen: Zehn Vorlesungen. VDM, Verlag Dr. Müller.

von Economo, C., Koskinas, G. N., & Triarhou, L. C. (1925). Atlas of cytoarchitectonics of the adult human cerebral cortex (Vol. 10). Karger Basel.

Wagstyl, K., Larocque, S., Cucurull, G., Lepage, C., Cohen, J. P., Bludau, S., Palomero-Gallagher, N., Lewis, L. B., Funck, T., Spitzer, H., Dickscheid, T., Fletcher, P. C., Romero, A., Zilles, K., Amunts, K., Bengio, Y., & Evans, A. C. (2020). BigBrain 3D atlas of cortical layers: Cortical and laminar thickness gradients diverge in sensory and motor cortices. PLOS Biology, 18(4), e3000678. https://doi.org/10.1371/journal.pbio.3000678

Zhang, J., Scholtens, L. H., Wei, Y., van den Heuvel, M. P., Chanes, L., & Barrett, L. F. (2020). Topography Impacts Topology: Anatomically Central Areas Exhibit a “High-Level Connector” Profile in the Human Cortex. Cerebral Cortex, 30(3), 1357–1365. https://doi.org/10.1093/cercor/bhz171

Zikopoulos, B., García-Cabezas, M. Á., & Barbas, H. (2018). Parallel trends in cortical gray and white matter architecture and connections in primates allow fine study of pathways in humans and reveal network disruptions in autism. PLoS Biology, 16(2), e2004559. https://doi.org/10.1371/journal.pbio.2004559

